# CRISPR-Cas9 target-strand nicking provides phage resistance by inhibiting replication

**DOI:** 10.1101/2024.09.05.611540

**Authors:** Giang T. Nguyen, Michael A. Schelling, Dipali G. Sashital

## Abstract

Cas endonucleases, like Cas9 and Cas12a, are RNA-guided immune effectors that provide bacterial defense against bacteriophages. Cas endonucleases rely on divalent metal ions for their enzymatic activities and to facilitate conformational changes that are required for specific recognition and cleavage of target DNA. While Cas endonucleases typically produce double-strand breaks (DSBs) in DNA targets, reduced, physiologically relevant Mg^2+^ concentrations and target mismatches can result in incomplete second-strand cleavage, resulting in the production of a nicked DNA. It remains poorly understood whether nicking by Cas endonucleases is sufficient to provide protection against phage. To address this, we tested phage protection by Cas9 nickases, in which only one of two nuclease domains is catalytically active. By testing a large panel of guide RNAs, we find that target strand nicking can be sufficient to provide immunity, while non-target nicking does not provide any additional protection beyond Cas9 binding. Target-strand nicking inhibits phage replication and can reduce the susceptibility of Cas9 to viral escape when targeting non-essential regions of the genome. Cleavage of the non-target strand by the RuvC domain is strongly impaired at low Mg^2+^ concentrations. As a result, fluctuations in the concentration of other biomolecules that can compete for binding of free Mg^2+^ strongly influences the ability of Cas9 to form a DSB at targeted sites. Overall, our results suggest that Cas9 may only nick DNA during CRISPR-mediated immunity, especially under conditions of low Mg^2+^ availability in cells.

## Introduction

CRISPR-Cas (Clustered regularly interspaced short palindromic repeats-CRISPR associated) systems provide sequence-specific protection to bacteria and archaea against viral infection (1–4). CRISPR arrays encode CRISPR RNAs (crRNAs) containing “spacer” sequences with complementarity to viral genomes (5–7). Cas effectors use these crRNAs as guides to find and destroy complementary nucleic acids upon infection (2, 3). New spacers can be acquired into the CRISPR array allowing for adaptation against specific viruses (1).

Cas effectors vary widely in their composition and mechanisms of action (8). Cas9 and Cas12a effectors target double-stranded DNA (dsDNA) and are composed of a single Cas endonuclease (9–11). Targeting dsDNA requires a protospacer adjacent motif (PAM) that is located next to the spacer-complementary target region (9–13). Cas endonucleases initially recognize PAM sequences and directionally unwind DNA away from the PAM, allowing for formation of an RNA-DNA hybrid between the crRNA spacer and the target strand of the DNA (14, 15). Formation of this R-loop enables cleavage by Cas endonuclease domains (16, 17). Cas9 contains two metal-dependent nuclease domains, HNH and RuvC, allowing simultaneous cleavage of both DNA strands to generate a double stranded break (DSB) (9, 10, 16). The HNH domain cleaves the crRNA-complementary “target” strand using a one-metal-ion-dependent mechanism, while the RuvC domain cleaves the opposite “non-target strand” via a two-metal-ion-dependent mechanism (9, 18). Active site mutations in either of these domains converts Cas9 into a nickase, which cleaves only a single strand in the DNA target (9, 10). Cas12a only contains a single RuvC nuclease domain, which is used to cleave the non-target and target strand in a sequential manner (19, 20).

In addition to serving as catalytic cofactors, metal ions are required for additional steps in the overall mechanisms of Cas9 and Cas12a. Mg^2+^ ions affect binding of both the guide RNA and the target DNA to Cas endonucleases (21–25). The Cas9 HNH domain requires divalent cations to transition from an inactive to an active conformation following target binding (26, 27). RuvC cleavage is modulated allosterically by these conformational changes of the HNH domain, but not by HNH nuclease function (16). Cas12a conformational changes also depend on metal ions. Mg^2+^-mediated local DNA unwinding in the PAM-distal region is required prior to the second, target-strand cleavage event by Cas12a (28).

Importantly, the physiological concentration of free Mg^2+^ is 1-2 mM in bacteria (29, 30). Previous studies have suggested that Cas endonucleases have defects in second strand cleavage at physiologically relevant Mg^2+^ concentration and especially upon introduction of PAM-distal mutations in the target DNA (24, 28, 31–35), which can occur in native settings when viral targets mutate under pressure of CRISPR-Cas immunity (13, 36–41). However, single point mutations in the PAM-distal region are rarely sufficient to allow complete escape from Cas9 or Cas12a. Together, these observations raise the question of whether nicking by Cas endonucleases is sufficient to allow anti-viral protection in cells.

To address this question, we compared the efficacy of wild-type (WT) Cas9 and Cas9 nickases in providing anti-bacteriophage protection in *Escherichia coli*. We accumulated a comprehensive dataset using >60 guide RNAs targeting either strand of the genome in coding and non-coding regions with varying essentiality. We find that nicking by the HNH domain, but not the RuvC domain, can provide protection at a similar level to WT Cas9, although the degree of protection by both WT Cas9 and the HNH nickase varies depending on target location. Phage proliferation is inhibited by both WT Cas9 and the HNH nickase due to decreased replication, while the RuvC nickase has little effect on the amount of phage DNA that accumulates over time. Using in vitro biochemistry, we show that the RuvC nickase, but not the HNH nickase, can be readily turned over by RNA polymerase. In addition, the RuvC nickase is highly susceptible to variable Mg^2+^ concentrations that may be altered due to fluctuations in nucleotide triphosphate (NTP) levels during phage infection. Overall, our results suggest that the low Mg^2+^ concentrations present in cells may result in nicking of target DNA by Cas9, and that increased free Mg^2+^ as NTPs are depleted may improve Cas9 activity during phage infection.

## Results

### Nicking by the Cas9 HNH domain is sufficient to provide anti-phage defense for some targets

We first investigated whether Cas9 nickases are sufficient to provide phage resistance. We introduced two plasmids for inducible expression of *Streptococcus pyogenes* (Sp)Cas9 and a single-guide RNA (sgRNA) in *E. coli* (Fig. 1A). These strains were infected with phage λvir, a mutant of phage λ that is locked into the lytic cycle (42). We tested the degree of anti-phage resistance for four SpCas9 variants: wild-type (WT) Cas9 that can produce DSBs; catalytically dead Cas9 (dCas9, D10A/H840A) that can bind but not cleave targets (43–45); an HNH nickase (D10A) that cleaves only the target strand; and a RuvC nickase (H840A) that cleaves only the non-target strand (9). We designed 61 sgRNAs to target various essential and non-essential regions of the lambda phage genome (Table S1). Targets were selected on both the template and coding strand, with several sgRNAs targeting the same or overlapping regions on either strand.

**Figure 1:**
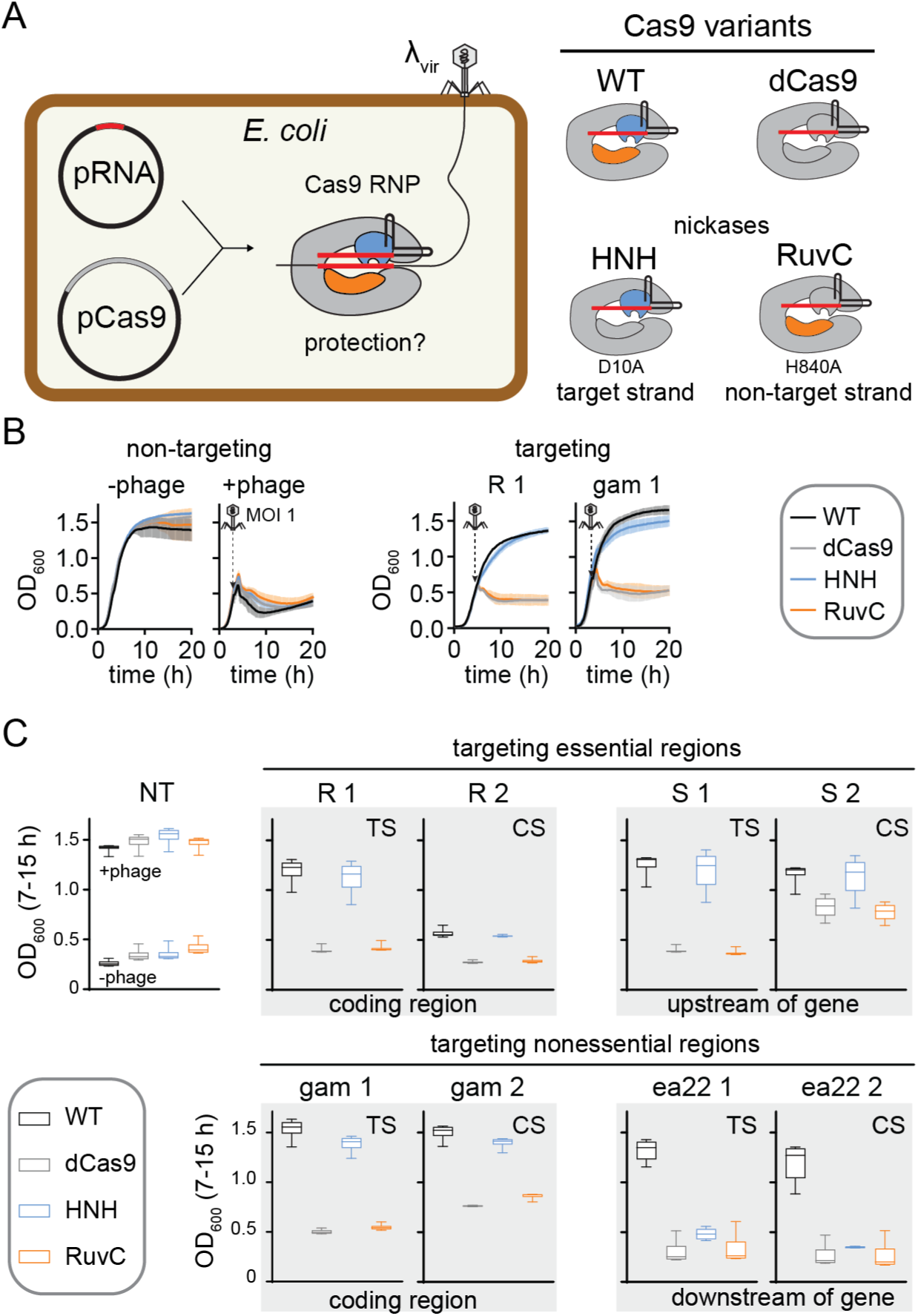
**Nicking by the Cas9 HNH domain can provide resistance against phage** A) Schematic of experimental design. Guide RNAs targeting various regions of the λvir genome were expressed from the pRNA plasmid and Cas9 variants shown on the right were expressed from pCas9. B) Representative growth curves for *E. coli* BW25113 expressing pRNA and pCas9 in the without (-phage) or with (+phage) of λvir infection at an MOI of 1. The growth curves on the left used a strain expressing a guide RNA that did not target the phage genome. The growth curves on the right expressed guide RNAs expressing gene R or *gam* in the λ genome. The average of three replicates are plotted with error bars representing standard deviation. C) Box plots of OD600 values over the 7-15 h time period of growth curves, summarizing protection by four sets of guide RNAs targeting the λ genome. Error bars show the minimum and maximum values. Plots that are outlined together in gray represent guide RNAs that target the same or overlapping regions on either the template strand (TS) or coding strand (CS). Additional targets are shown in Figure S1.

The growth of infected liquid cultures expressing Cas9 variants and sgRNA was monitored over time by measuring OD600 every 10 min (Fig. 1B). To easily compare protection by each variant, we plotted OD600 values at each time point from 7-15 h from three replicate cultures (Fig. 1C, S1, 2A). During this time range, cultures in which Cas9 provides protection have a similar OD600 to uninfected cells, while cultures that have lysed have a substantially lower OD600 (Fig. 1C).

**Figure 2:**
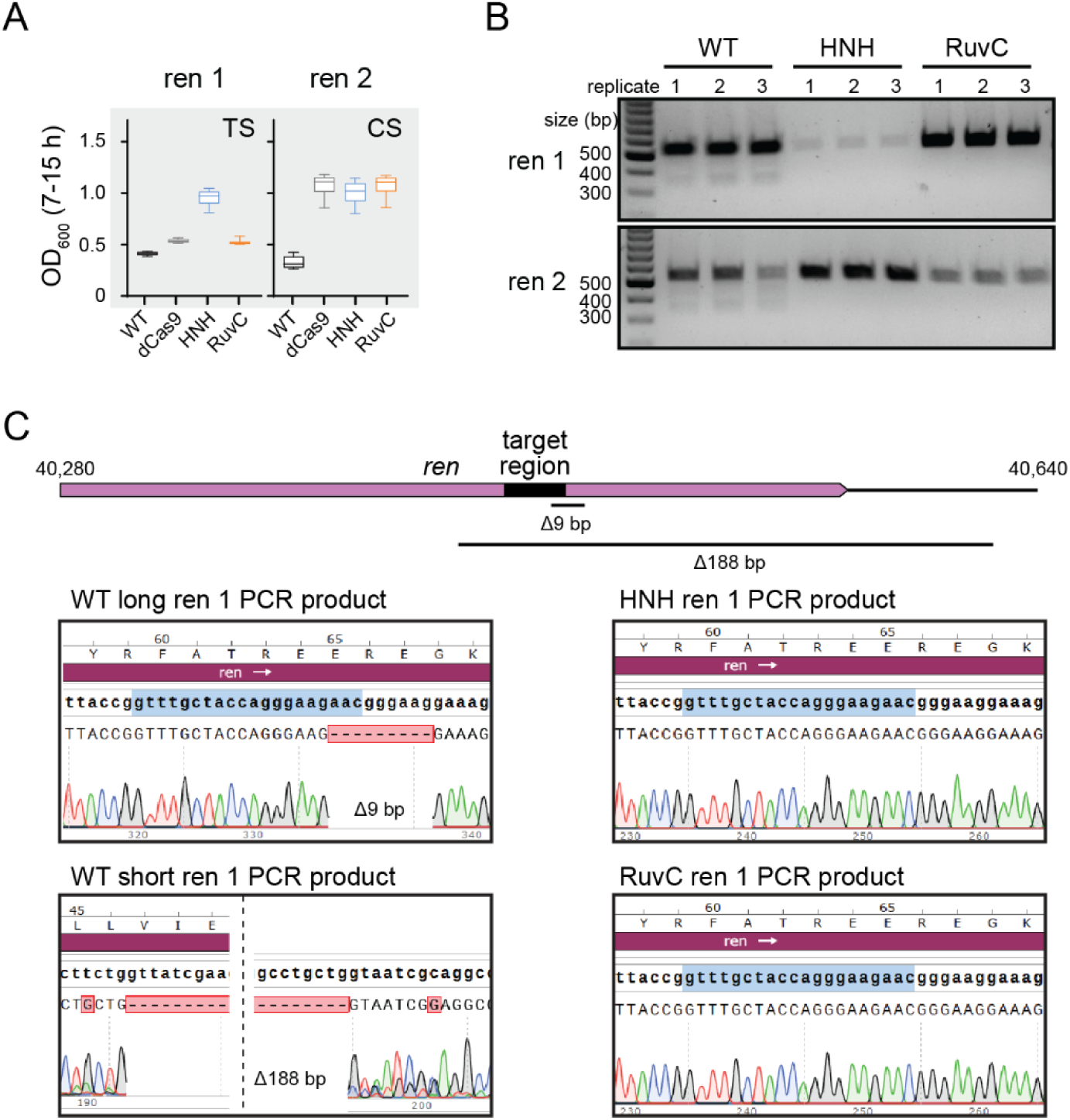
Nicking provides protection and prevents deletion of target region A) Box plots (as in Fig. 1C) for *E. coli* cells expressing Cas9 variants and one of two guide RNAs targeting the same region on the template (TS) or coding (CS) strand of the *ren* gene, a non-essential gene in the λ genome. B) Agarose gel for PCR amplicons of the *ren* gene from phage recovered following growth of cultures plotted in (A). C) Sanger sequencing chromatograms for *ren* PCR amplicons for cultures expressing WT Cas9, HNH nickase, or RuvC nickase. PCR amplicons of ∼550 bp were sequenced for the long WT Cas9 PCR product and both nickase products. The short WT Cas9 PCR was purified from the band at ∼350 bp on the gel in (B).

We observed strong protection by WT Cas9 for most targets (Fig. 1C, S1). It has previously been reported that guide RNAs targeting the template strand of phage targets provide stronger protection than those targeting the coding strand (41, 46). However, we did not observe any consistent trends in the relative amount of protection by WT Cas9 when targeting overlapping regions on the template versus the coding strand. We did not observe substantial protection by any Cas9 variants for a few targets (e.g. nu1, ea31, Fig. S1), which may be due to misfolding of the sgRNA or high G/C content within the targeted region.

The amount of protection provided by dCas9 and the two Cas9 nickases varied significantly. For most targets, we observed no or low protection by dCas9, suggesting that transcriptional repression by Cas9 binding^38–41, 43^ is usually not sufficient to provide protection. While we generally observed a similar lack of protection by the RuvC nickase, we consistently observed more protection by the HNH nickase. We observed similar protection by WT Cas9 and the HNH nickase for several targets and partial protection by the HNH nickase for most other targets (Fig. 1C, S1). Overall, these results suggest that nicking by Cas9 is sufficient to provide at least partial protection against phage infection, but that nicking must occur by the HNH rather than the RuvC domain.

### Genomic deletions allow phage escape from WT Cas9 but not nickases

Surprisingly, for a few targets, we observed better protection by the HNH nickase than WT Cas9, especially when targeting non-essential regions of the lambda phage genome. In particular, two sgRNAs targeting overlapping regions of the *ren* gene on either the template or coding strand provided no protection when paired with WT Cas9 and partial protection by the HNH nickase or by all three other Cas9 variants (Fig. 2A). It has previously been observed that Cas effector cleavage can result in mutagenesis within the target region of phage genomes, including point or deletion mutations (36, 39–41). Such mutations could allow phages to escape Cas9-mediated immunity, resulting in the lack of protection by WT Cas9.

To determine whether Cas9 cleavage resulted in target mutations, we PCR amplified phage genomic regions containing the *ren* targets following challenge with WT Cas9 or either Cas9 nickase. We observed a single product of the expected size for phages challenged with the nickases, but an additional light smear of products smaller than the expected size when WT Cas9 was used (Fig. 2B). Sequencing of the PCR amplicons revealed that phages challenged with WT Cas9 contained a mixture of different deletions in the targeted region. The full-length amplicon contained a short 9-bp deletion that removed the PAM and first three nucleotides of the seed. The short PCR amplicon contained a larger 188-bp deletion that removed the entire target site. In contrast, both Cas9 nickases retained the original target sequence. Overall, these results suggest that nicking by Cas9 may allow enhanced protection by reducing the tendency of phages to develop escape mutations following Cas9 targeting.

### Target strand nicking blocks phage replication

We next asked how target strand nicking by the HNH nickase provides protection. Nicking has previously been shown to inhibit phage genome replication (47). To test whether nicking by Cas9 nickases similarly blocks phage replication, we used quantitative PCR (qPCR) to measure phage DNA replication efficiency upon challenge by each Cas9 variant. We selected two guide RNAs (R1 and Q1) that conferred protection only when paired with WT Cas9 or the HNH nickase (Fig. 3A). To test phage replication, mid-log cultures were infected with λvir at an MOI of 0.2 for 10 min (Fig. 3B). The cells were then pelleted, washed three times, and grown in fresh media. An aliquot of the cultures was harvested immediately after adsorption and following 30 or 60 min of growth. Cells infected at an MOI of 0.2 did not lyse within 60 min (Fig. 3A), indicating that the selected time-points occur within the timeframe of a single infection cycle.

**Figure 3:**
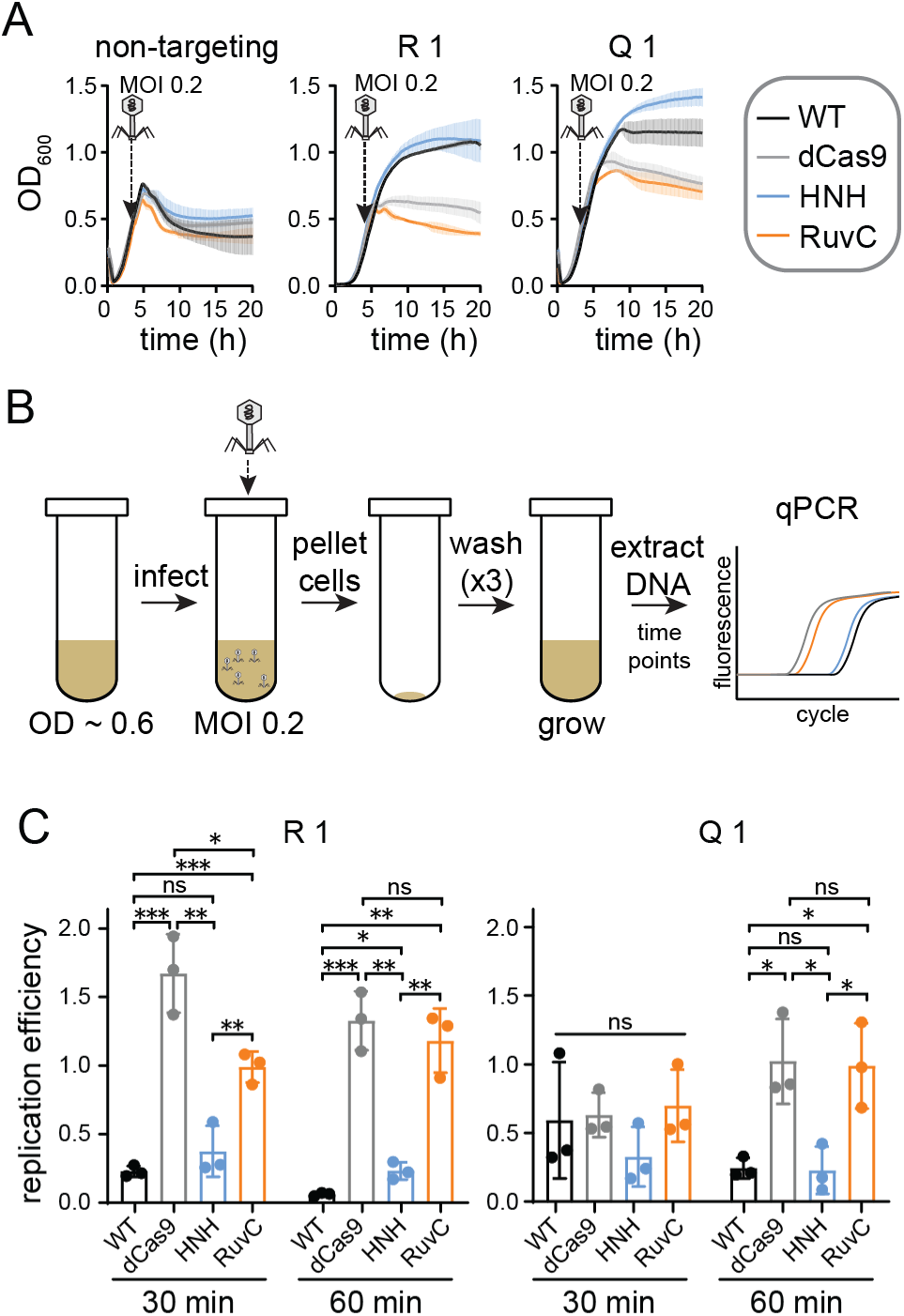
HNH domain nicking reduces phage DNA replication A) Growth curves for cultures used for qPCR analysis. *E. coli* BW25113 expressing a Cas9 variant and a non-targeting control guide RNA, or guide RNAs targeting gene R or Q in the λ genome, were infected with λvir at an MOI of 0.2. B) Schematic of phage infection to ensure a single infection cycle. *E. coli* BW25113 expressing Cas9 variants and guides used in (A) were grown to mid-log phase. Phage was added at an MOI of 0.2 and incubated for 10 min to allow adsorption, then cells were pelleted and washed with fresh LB three times. Cells were grown for 60 min, and phage DNA was assayed using qPCR at 30 and 60 min. C) Replication efficiency of λvir in cultures expressing each Cas9 variant. Replication efficiency was calculated by normalizing Ct values for cultures expressing a targeting guide RNA versus cultures expressing the non-targeting guide (see Methods). The average value of three replicates is shown, with individual data points shown as dots. Error bars represent standard deviation. *P* values were determined using an unpaired, two-tailed t-test. ns: *P* > 0.05, *: *P* < 0.05, **: *P* < 0.01, ***: *P* < 0.001.

The harvested samples were boiled to prevent further DNA replication, then were used as DNA templates for qPCR (see Methods).

Relative to cultures expressing a non-targeting guide, phage DNA decreased significantly in cells expressing WT Cas9 and either the R1 or Q1 guide (Fig. 3C). We observed similarly low relative amounts of phage DNA from cultures expressing the HNH nickase and either guide.

Phage DNA was significantly higher for the dCas9 and RuvC cultures at both times points for the R1 guide, but only at the 60 min timepoint for the Q1 guide, suggesting that Cas9 binding may have initially inhibited phage replication in cultures expressing the Q1 guide. Consistently, we observed a longer period of growth following addition of phage to liquid cultures expressing dCas9 and the RuvC nickase bearing the Q1 guide than for cultures with a non-targeting guide (Fig. 3A). Overall, these results suggest that nicking by the HNH domain is sufficient to inhibit replication of phage DNA, but that Cas9 binding or nicking by the RuvC domain do not substantially impair phage replication over the course of an infection cycle.

### Non-target strand nicking is necessary for Cas9 turnover by RNA polymerase

Our phage assays suggest that nicking of the target strand by the HNH domain can provide protection from λvir on par with WT Cas9, but that nicking of the non-target strand by the RuvC domain provides no additional protection beyond Cas9 binding. One potential explanation for the difference between target and non-target strand nicking is the ability for Cas9 to turn over following target cleavage. Although Cas9 typically remains bound to its product following cleavage, it has previously been demonstrated that RNA polymerase can dislodge Cas9 from the template strand of a DNA substrate (46). Conversion of Cas9 into a multi-turnover enzyme has been suggested to improve protection against phage. Thus, it is possible that differences between turnover of HNH and RuvC nickases may account for the differences in protection observed for these Cas9 variants.

To test this, we performed Cas9 cleavage assays in the absence and presence of T7 RNA polymerase (T7 RNAP) (Fig. 4A). The substrate plasmid contained a T7 promoter upstream of the target site, which was located either on the template or the coding strand. We performed cleavage assays using an excess of the plasmid over Cas9-guide RNA complex. In the absence of T7 RNAP, all three Cas9 variants cleaved 5-20 percent of the target DNA, consistent with an inability of the enzymes to turnover following a single round of cleavage (Fig. 4B-C, S2A-B).

**Figure 4:**
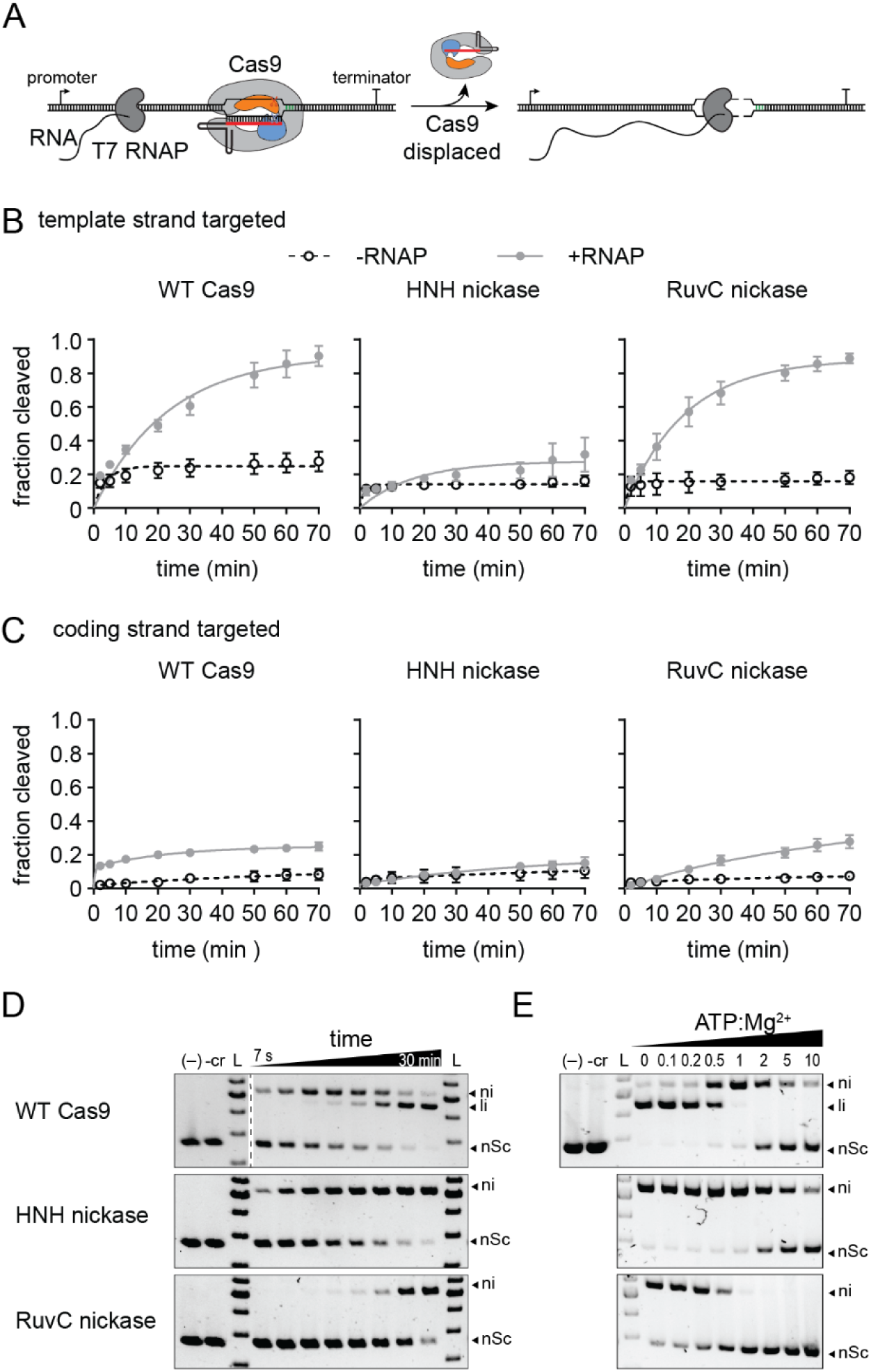
Target strand nicking prevents Cas9 turnover and occurs under physiological metal ion conditions A) Schematic of Cas9 turnover assay. The Cas9 target was located on the template strand downstream of a T7 RNA polymerase promoter. WT Cas9 typically remains bound to the target following cleavage, but can be dislodged by RNA polymerase transcription of the target region. B-C) Fraction cleaved over time for the (B) template or (C) coding strand by WT Cas9, HNH nickase, or RuvC nickase in the absence (open circles, dashed lines) or presence (closed gray circles, solid line) of RNA polymerase. The average of three replicates are shown and error bars represent standard deviation. Representative gels are shown in Fig. S2. D) Target plasmid cleavage time course for WT Cas9, HNH nickase, and RuvC nickase at 1 mM MgCl2 concentration. The two controls contain only plasmid (-) or Cas9 without guide RNA (-cr) incubated for the longest time point. Nicked (ni), linear (li) and negatively supercoiled DNA species are labeled. Time points: 7 s, 15 s, 30 s, 1 min, 2 min, 5 min, 15 min, and 30 min. E) Target plasmid cleavage at varying ATP:Mg^2+^ ratios. The MgCl2 concentration was 1 mM and each reaction was incubated for 30 min.

**Figure 5:**
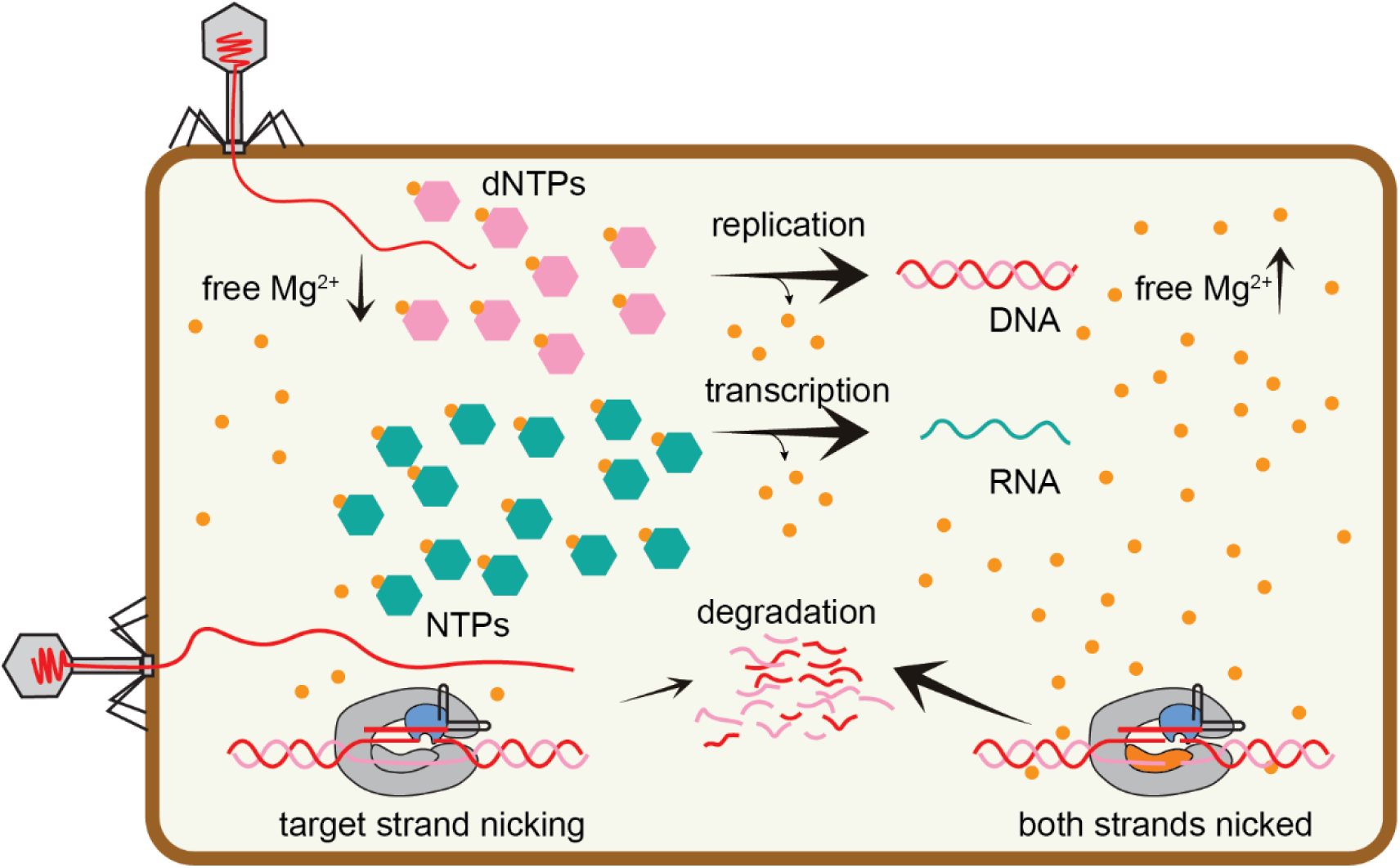
Metal-dependent nicking during Cas9-mediated phage defense Under typical cellular conditions, cleavage of the non-target strand by the RuvC domain may be impaired due to low Mg^2+^ availability. Target strand nicking by the HNH domain may be sufficient to provide protection during early stages of phage infection. However, if phage infection proceeds, replication and transcription of the phage genome results in depletion of nucleotides, freeing Mg^2+^ and allowing increased cleavage of both strands by Cas9.

The fraction cleaved by WT Cas9 increased substantially upon addition of T7 RNAP when targeting the template strand, indicating that Cas9 is dislodged by T7 RNAP allowing for further cleavage of the target (Fig. 4B, S2A). Surprisingly, we observed a similar turnover rate for the RuvC, but not the HNH nickase, suggesting that nicking of the non-target strand is necessary to allow Cas9 turnover by RNAP. We similarly observed more turnover for WT Cas9 and the RuvC nickase when targeting the coding strand, albeit to a much smaller degree than for the template strand (Fig. 4C, S2B). It has previously been observed that RuvC cleavage exposes the 3’ end of the non-target strand, while the target strand remains hybridized with the crRNA after cleavage by the HNH domain (48). Together with our turnover results, these observations suggest that the exposed nicked non-target strand is necessary for RNA polymerase to dislodge Cas9 from target DNA following cleavage by the RuvC domain.

### Non-target strand cleavage is slow at physiological metal ion conditions

Our results suggest that the rate of Cas9 turnover does not account for the differences in protection provided by the two Cas9 nickases. An alternative explanation may be the degree of cleavage by each nickase that may occur under physiological conditions. It has previously observed that the RuvC domain requires two Mg^2+^ ions and may bind to metal with a lower affinity than the HNH domain (18). We observed rapid nicking but slow linearization of plasmid DNA by WT Cas9 at 1 mM Mg^2+^, suggesting that the two nuclease domains cleaved each strand of the DNA at different rates (Fig. 4D). Cleavage by the HNH nickase was more consistent with the rate of nicking by WT Cas9, while the rate of nicking by the RuvC nickase was similar to the rate of linearization of the plasmid by WT Cas9. These results suggest that slow cleavage by the RuvC nickase limits its ability to provide anti-phage protection under low, physiological metal ion conditions.

Cellular Mg^2+^ concentrations are estimated in the 10-100 mM range, but much of this is expected to be bound by various biomolecules in cells (49, 50). For example, nucleotide triphosphates (NTPs) can chelate metal ions with a *K*d in the sub-mM range (51). Thus, enzymes like Cas9 that are highly sensitive to metal ion concentrations may also be sensitive to concentrations of other metal-binding biomolecules in cells. To simulate this, we performed plasmid cleavage assays in the presence of 1 mM Mg^2+^ and increasing concentrations of ATP (Fig. 4E). We observed a substantial loss of cleavage by the RuvC domain at sub-stoichiometric amounts of ATP:Mg^2+^, and a complete loss of cleavage at equimolar amounts of ATP and Mg^2+^. In contrast, the HNH domain was able to cleave the DNA until the ATP concentration exceeded Mg^2+^. Together, these results suggest that the RuvC domain of Cas9 is highly sensitive to metal ion conditions as well as to the concentration of other biomolecules that may fluctuate during phage infection.

## Discussion

DNA-targeting Cas effectors can provide robust defense against phage by directly damaging viral DNA. Canonically, Cas9 and Cas12 effectors are thought to induce double-strand breaks that enable degradation of the invading genome through the activity of other cellular nucleases (9–11). However, both Cas9 and Cas12a are prone to nicking of DNA at low, physiologically relevant metal ion concentrations or in the presence of target mismatches (24, 28, 31–35). Our study reveals that nicking by Cas endonucleases can be sufficient to enable immunity. We observed stronger protection when the Cas9 HNH domain nicked the target strand, often on the same order as for WT Cas9. Nicking may be beneficial during phage infection because it can reduce the rate of phage replication while decreasing the likelihood of escape mutations arising following DSB formation at the target site. Indeed, we observed that nicking by the HNH domain greatly reduced the amount of phage DNA present during an infection cycle and also prevented deletion mutations that otherwise arose upon targeting of non-essential genomic regions by WT Cas9.

It has previously been observed that Cas9 provides better protection when targeting the template strand than the coding strand, potentially due to the turnover of Cas9 by RNA polymerase (40, 46). However, when testing a larger pool of guide RNAs targeting overlapping regions on the template and coding strand, we did not observe a consistent strand bias in Cas9- mediated immunity. Furthermore, the HNH nickase often exhibited efficient phage resistance, although we did not observe turnover of the HNH nickase by RNA polymerase. In contrast, the RuvC nickase, although dislodged by RNA polymerase, did not provide protection in our phage assays. These results suggest that Cas9 turnover may not contribute to the efficiency of phage resistance. It is also possible removal of the RuvC nickase allowed rapid repair of the nicked DNA, compromising the effectiveness of the RuvC nickase in combating phage infections. Similarly, it has been proposed that turnover of the H840A RuvC nickase by the replication fork remodeler HLTF allows rapid repair of nicks, preventing effective genome editing by the RuvC nickase in human cells (52).

Although the RuvC and HNH domains are thought to act synergistically (16), recent studies have suggested that cleavage of the target strand by the HNH domain may occur prior to cleavage of the non-target strand by the RuvC domain (25, 35). Our *in vitro* assays also showed that cleavage of the non-target strand by the RuvC domain is substantially slower than for the HNH domain at low, physiologically relevant Mg^2+^ concentrations. RuvC cleavage is also impaired by the presence of other biomolecules, like ATP, that can compete for binding to Mg^2+^. The increased sensitivity of RuvC to metal ion concentrations may be due to its requirement for two Mg^2+^ ions, in comparison to only one for the HNH domain, or to reduced affinity for metal ions within the RuvC active site (18).

Together, our data suggest a model in which Cas9 may only nick DNA under conditions where free Mg^2+^ is limited in cells. Nicking may be sufficient to reduce phage replication while also preventing deletions within targeted non-essential genomic regions. Thus, nicking may be beneficial to preserve the effectiveness of spacer sequences for longer periods. Increased transcription and replication during a phage infection cycle depletes nucleotides, likely resulting in release of free Mg^2+^. While Cas9 is thought to target phage early in the infection cycle (53), increased Cas9 cleavage activity upon release of Mg^2+^ could act as a safeguard to ensure protection during the late stages of infection. This type of regulatory mechanism may be especially important to ensure continued cleavage upon introduction of point mutations within the targeted region (39, 40).

Other Cas endonucleases, including Cas12a, have also been observed to nick DNA in a metal-dependent manner (24, 36). We previously observed that mutations arise preferentially in the PAM-distal region upon targeting by Cas12a (36). PAM-distal mutations cause substantial target-strand cleavage defects at lower Mg^2+^, which are alleviated at higher metal ion concentrations (24). While it remains unclear whether nicking by Cas12a is sufficient to provide protection against phage, it is notable that Cas12a nicks the non-target, rather than the target strand. Thus, it is possible that Cas12a is more susceptible to escape via PAM-distal mutations than Cas9, as we have previously observed (36).

## Materials and Methods

### Expression plasmid construction

All plasmids and primers used in this study are listed in the Supplementary tables. SpCas9 expression plasmids and crRNA expression plasmids were constructed using pACYCDuet-1 and pUC19, respectively. Both Cas9 and crRNA were inserted downstream of pBAD promoter as previously described (36). Mutations were introduced into the Cas9 construct by round-the-horn PCR and ligation to create the HNH (D10A) or RuvC (H840A) nickases. A second mutation was introduced to produce the dCas9 expression construct (D10A, H840A).

For Cas9 purification, the expression vector pMJ806 was a gift from Jennifer Doudna (Addgene plasmid # 39312 ; http://n2t.net/addgene:39312 ; RRID:Addgene_39312) (9). The D10A or H840A substitutions were introduced via round-the-horn for expression of the HNH and RuvC nickase, respectively.

### Liquid culture phage assays and growth curves

The phage challenge assays were conducted using *E. coli* strain BW25113 which is a derivative of K12 (54). A virulent mutant of phage Lambda (λvir) that is locked lysogenic cycle to create a purely lytic phage (42). Cas9 and crRNA expression plasmids were introduced into *E. coli* by heat shock. The transformed cells were plated on LB agar plates containing ampicillin (100 μg/mL) and chloramphenicol (25 μg/mL) for selection. A single colony was cultured overnight in LB media with the same antibiotics as above. Fresh 200 µL LB cultures supplemented with 100 μg/mL ampicillin, 25 μg/mL chloramphenicol, 20 mM arabinose, and 10 mM MgSO4 were inoculated with the overnight cultures in a 1:100 ratio. Lambda phage was added at MOI 1 at OD600 of 0.6. The cultures were grown in a TECAN infinite M Nano+ 96-well plate reader at 280 rpm and 37 °C and OD measurements at 600 nm wavelength were measured every 10 min. All targets for phage assays are listed in the Supplementary tables.

### WT Cas9 and Cas9 nickase protein expression and purification

The expression of Cas9 proteins was carried out in *E. coli* BL21(DE3) cells. To initiate the expression, overnight culture of the cells carrying the respective expression plasmid was added into fresh LB media (at a 1:100 ratio) supplemented with 50 μg/mL kanamycin. When the culture reached an OD600 of 0.5 to 0.6, protein expression was induced by the addition of 0.5 mM IPTG. The induced culture was then incubated at 18°C for approximately 16 hours with continuous shaking. Cells were centrifuged at 6000 ×g for 15 min at 4 °C.

Harvested cells were resuspended in lysis buffer containing 20 mM Tris-HCl (pH 8.0), 500 mM NaCl, 5 mM imidazole, and 5% glycerol. 1 mM PMSF was added before sonication and the lysate was centrifuged to remove insoluble material. The clarified supernatant was transferred to a HisPur Ni-NTA resin column (Thermo Fisher Scientific) pre-equilibrated with lysis buffer.

The column was washed with 50 column volumes of lysis buffer, followed by a second wash with 50 column volumes of wash buffer (20 mM Tris-HCl (pH 8.0), 500 mM NaCl, 15 mM imidazole, and 5% glycerol). An elution buffer composed of 20 mM Tris-HCl (pH 8.0), 500 mM NaCl, 250 mM imidazole, and 5% glycerol was applied. The eluted protein fractions were collected and analyzed by SDS-PAGE. Fractions containing MBP-Cas9 were combined and cleaved with TEV protease in a 1:100 (w/w) ratio to remove the MBP tag at 4 °C while dialyzing against dialysis buffer (10 mM HEPES-KOH (pH 7.5), 200 mM KCl, 1 mM DTT, and 5% glycerol). After overnight dialysis, the protein was diluted with a buffer containing 20 mM HEPES-KOH (pH 7.5) and 5% glycerol to a final concentration of 100 mM KCl. The protein was loaded on a HiTrap Heparin HP (Cytiva) column which was pre-equilibrated with Buffer A (20 mM HEPES-KOH (pH 7.5), 100 mM KCl, and 5% glycerol). 20% of buffer B (20 mM HEPES-KOH (pH 7.5), 1 M KCl, and 5% glycerol) was applied to wash the column. A gradient from 20% to 100% of buffer B was applied over a total volume of 60 mL to elute. Protein fractions were collected and analyzed using SDS-PAGE. Fractions containing the protein of interest were pooled and concentrated to a final volume of 1 mL. The concentrated protein was diluted to 15 mL with storage buffer (20 mM HEPES-KOH, pH 7.5, 200 mM KCl, and 1 mM DTT). This buffer exchange step was performed three times. Finally, the concentrated protein was aliquoted, flash-frozen in liquid nitrogen, and stored at −80 °C for future use.

### T7RNA polymerase expression and purification

T7 RNA polymerase expression and purification was performed by the following protocol. pTT7-911Q (T7 RNA polymerase expression plasmid) was transformed into BL21(DE3) and plated on LB containing 50 µg/ml ampicillin. A single colony was picked and grown overnight in 25 mL LB medium supplemented with 50 µg/ml ampicillin. To initiate the expression, the overnight culture was added into fresh LB media (at a 1:100 ratio) supplemented with 50 µg/ml ampicillin and grown at 37 °C to OD600 of 0.5 to 0.6. Protein expression was induced by adding 0.5 mM ITPG and then was continued to grow at 37 °C for 3 hours. The cell pellets were harvested at 6000 xg for 15 min at 4 °C.

The cell pellets were resuspended in lysis buffer containing 50 mM Tris-HCl (pH 8.0), 100 mM NaCl, 5 mM β-Mercaptoethanol, 1 mM imidazole, and 5% glycerol was used to resuspend the cell pellet. 0.1 mM PMSF was added into the resuspended cells right before by sonication step and the cell lysate then was centrifuged at 20, 000 ×g for 30 min. The lysate was incubated with 2 mL HisPur resin (pre-equilibrated with lysis buffer) with gentle rocking for 1 hour at 4 °C. The resin was pellet at 1000 xg for 2 min, then the lysate was carefully poured off. The resin was washed 4 times with lysis buffer and then 4 times with wash buffer (50 mM Tris pH 8, 100 mM NaCl, 5 mM β-Mercaptoethanol, 10 mM imidazole, and 5% glycerol) using batch method. The protein was eluted using elution buffer (50 mM Tris pH 8, 100 mM NaCl, 5 mM β-Mercaptoethanol, 200 mM imidazole, and 5% glycerol). After analyzing the collected fractions using SDS-PAGE gel, the fractions containing the protein of interest were pooled together and concentrated to a final volume of 1 mL. The concentrated protein was run on a Superdex 200 column (Cytiva) using SEC buffer (50 mM Tris pH 8, 100 mM NaCl, 5 mM β-Mercaptoethanol, and 5% glycerol). Protein fractions were analyzed and dialyzed overnight using dialysis buffer (20 mM Sodium phosphate pH 7.7, 100 mM NaCl, 50% glycerol and 1 mM DTT). Finally, the protein was aliquoted and stored at −20°C.

### crRNA and tracrRNA preparation

crRNA and tracrRNA were in vitro transcribed by the following protocol. For the crRNA, short oligonucleotides (Integrated DNA Technologies) consisting of a T7 promoter region, and a template sequence were initially pre-annealed to a complementary T7 promoter sequence by heating at 90 °C for 2 min, followed by incubation at room temperature for 10 min. The tracrRNA template was cloned into the pUC19 plasmid, incorporating an EcoRI restriction site at the end of the template sequence. The tracrRNA plasmid was initially linearized using EcoRI and subsequently used as a template for in vitro transcription without the pre-annealing step.

Transcription reactions (500 μL) were performed in transcription buffer containing 40 mM Tris (pH 8.0), 38 mM MgCl2, 1 mM Spermidine (pH 8.0), 0.01% Triton X-100, 5 mM ATP, 5 mM CTP, 5 mM GTP, 5 mM UTP, and 5 mM DTT. The reaction mixture contained 0.5 μM pre-annealed DNA or 35 μg plasmid DNA and 1 μM T7 RNA polymerase. The transcription reaction was performed at 37 °C for 4 hours. Following the reaction, 2X RNA dye (New England Biolabs) was added, and the mixture was heated at 95 °C for 5 minutes, and then immediately cooled on ice. The reaction mixture was separated on a 10% polyacrylamide gel containing 1X TBE and 8 M urea. The crRNA band was visualized under UV-light and excised from the gel.

The gel slice was crushed and soaked overnight in 1 mL of nuclease-free water at 4 °C with gentle agitation. After centrifuging the gel mixture for 5 min at 2, 000 × g, the crRNA solution was transferred to Costar Spin-X centrifuge tube filters (Sigma Aldrich) and then centrifuged at the highest speed for 2 min to collect the crRNA solution in the collection chamber. crRNA was concentrated by ethanol precipitation, aliquoted and stored at -20 °C.

### In vitro cleavage assays

Cleavage assays were performed as described in a previous study (36). The reaction buffer contained 20 mM HEPES (pH 7.5), 100 mM KCl, 1 mM or 10 mM MgCl2, 1 mM DTT, and 5% glycerol. The final concentration of SpCas9 was 50 nM. The RNP complex was formed by incubating Cas9, crRNA, and tracRNA at a ratio of 1:1.5:1.5 at 37 °C for 10 minutes. To initiate the cleavage reaction, pre-warmed target plasmid at 37 °C was added to a final concentration of 15 ng/μL and a final reaction volume of 100 μL. The reaction was continuously incubated at 37 °C and 10 μL of aliquots were quenched at 7, 15, 30, 60, 300, 900, and 1, 800 s by 10 μL phenol-chloroform-isoamyl alcohol (25:24:1 v/v, Invitrogen). Following extraction of the aqueous layer, the cleavage products were separated on a 1% agarose, stain free gel and visualized using SYBR Safe by post-staining the gel (Invitrogen).

For Cas9 turnover assays, the reaction buffer contained 20 mM HEPES (pH 7.5), 100 mM KCl, 10 mM MgCl2, 1 mM DTT, 1 mM rNTP and 5% glycerol. The Cas9 RNP was formed as described above. The concentration of Cas9 was 10 nM and the final concentration of target plasmid was 20 ng/μL (∼12 nM). 1 μM T7 RNA polymerase was added 1 min after initiating the reaction. 10 μL aliquots were quenched at 2, 5, 10, 20, 30, 50, 70 min and separated on a 1% agarose gel with post-staining using SYBR Safe. The intensity of individual DNA bands was quantified using ImageJ software (55). To determine the fraction of cleaved DNA, the total cleaved DNA (including nicked and linearized DNA) was divided by the total DNA (nicked, linearized, and supercoiled DNA). The fraction cleaved was plotted against time, and the data was fitted to a double-exponential equation in GraphPad Prism. Each measurement was performed in triplicate.

For cleavage assays performed in the presence of ATP, the reaction buffer contained 20 mM HEPES (pH 7.5), 100 mM KCl, 1 mM MgCl2, 1 mM DTT, various concentrations of ATP (0.1, 0.2, 0.5, 1, 2, 5, 10 mM) and 5% glycerol. The final concentration of SpCas9 was 50 nM and the RNP was formed as described above.

### qPCR sample preparation

The transformed cells were grown in LB medium supplemented with ampicillin (100 μg/mL) and chloramphenicol (25 μg/mL) to an OD600 of 0.5-0.6 and infected with lambda phage at MOI 0.2. The cultures were incubated at 37 °C for 10 min without shaking to allow phage to inject their genetic material into the host cells. The cultures were centrifuged at 5000 × g for 5 min. The supernatant was discarded, and the cell pellets were resuspended in fresh LB medium and pelleted again as above to remove excess phage. The cell pellets were resuspended in fresh LB. 1 ml of this culture was harvested immediately (T0 samples) and the remaining culture was incubated at 37 °C with shaking. At time = 30-, and 60-min post-infection, 1 mL samples of the culture were harvested. To stop replication, the collected samples were heated at 100 °C for 10 min. The samples were diluted 10X with nuclease free water and directly used as template for qPCR.

### Quantitative PCR

qPCR reactions (20 µL) contained 1X SYBR green mix (Thermo Fisher), 500 nM of phage-specific primers or genome-specific primers and 2 µL of prepared samples described above. QuantStudio3 Real-Time PCR system (Thermo Fisher) was used to amplify the DNA templates according to the following program: one cycle of 50°C for 2 minutes, one cycle of 95 °C for 10 min, 40 cycles of 95 °C for 15 seconds and 60 °C for 60 seconds. PCR reactions were performed using primers amplifying gene L in the λ genome or the glyceraldehyde-3-phosphate dehydrogenase (*gap*) gene from the *E. coli* genome (Supplementary tables).

DNA replication was determined by quantifying the fold change in replication at *T* = 30 or 60 compared to genome copy at *T* = 0. The Ct value of phage samples at 0 min post-infection was used to calibrate the remaining Ct values. The Ct values for the *gap* gene from the host *E. coli* genome was used for normalization. To determine the fold change, the equation from the Real-Time PCR Handbook (ThermoFisher) was used.

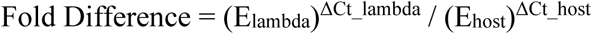

where

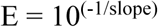

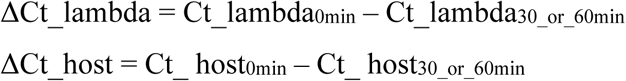

Replication efficiency was determined by normalizing the fold change for each sample to host cells expressing non-target sgRNA and similarly grown in the presence of antibiotics and inducer. Three biological replicates were performed for each sample.

## Acknowledgments

We would like to thank Dr. Hua Bai from Iowa State University, Department of Genetics, Development, and Cell Biology for technical and equipment support for qPCR experiment. We thank Akshara Raju and other members of the Sashital lab for helpful discussions. This work was supported by NSF grant 1652661 and NIH grant GM140876 (D.G.S.).

---

**Figure S1:**
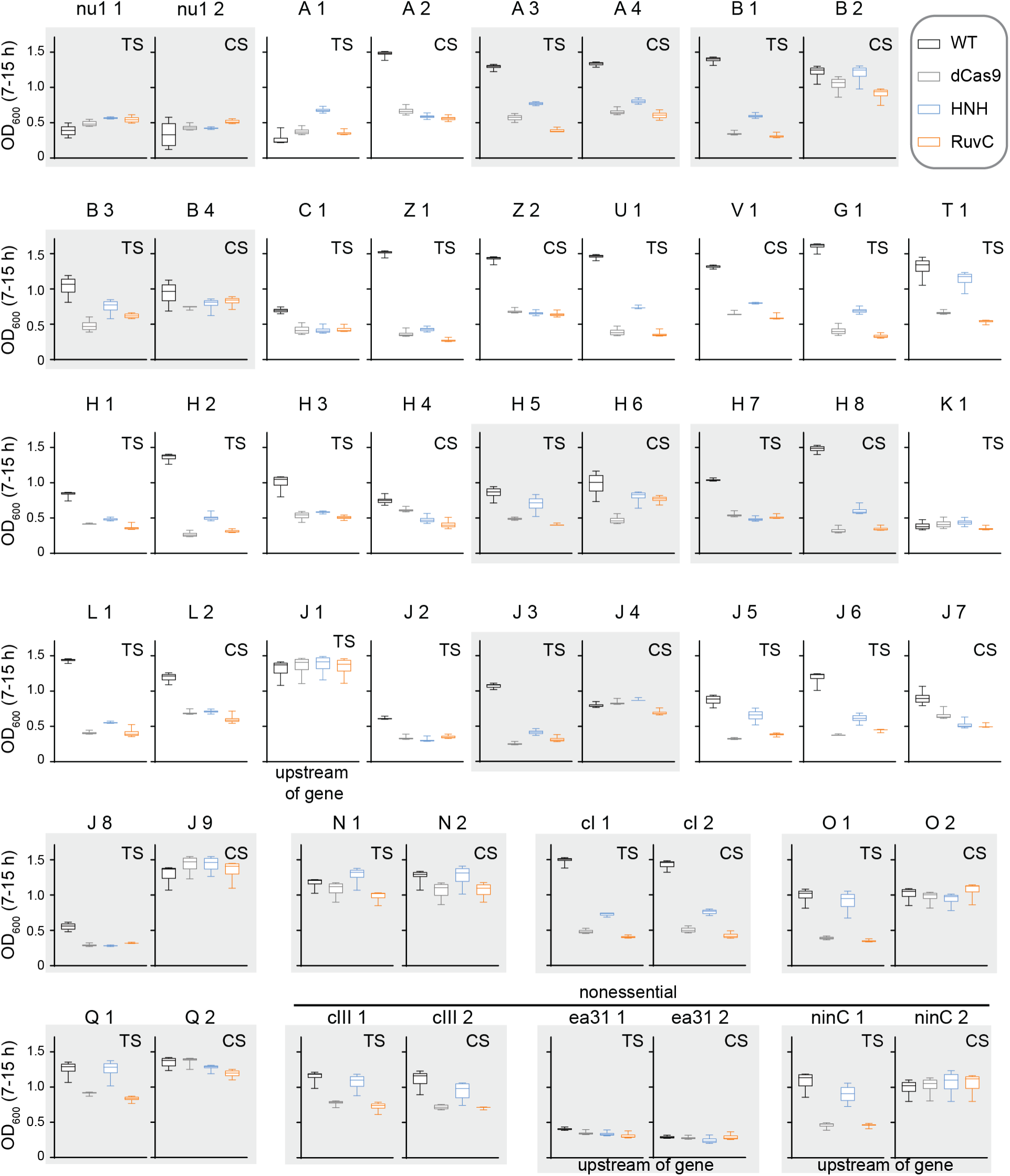
**Protection by Cas9 variants for additional guide RNAs.** Box plots of OD600 values over the 7-15 h time period of growth curves, summarizing protection by four Cas9 variants using different guide RNAs. The gene targeted by the guide RNA is indicated. Plots that are outlined together in gray represent guide RNAs that target the same or overlapping regions on either the template strand (TS) or coding strand (CS). Guide RNAs targeting non-essential genes or non-coding regions are indicated. All gene targets are provided in the Supplementary tables.

**Figure S2:**
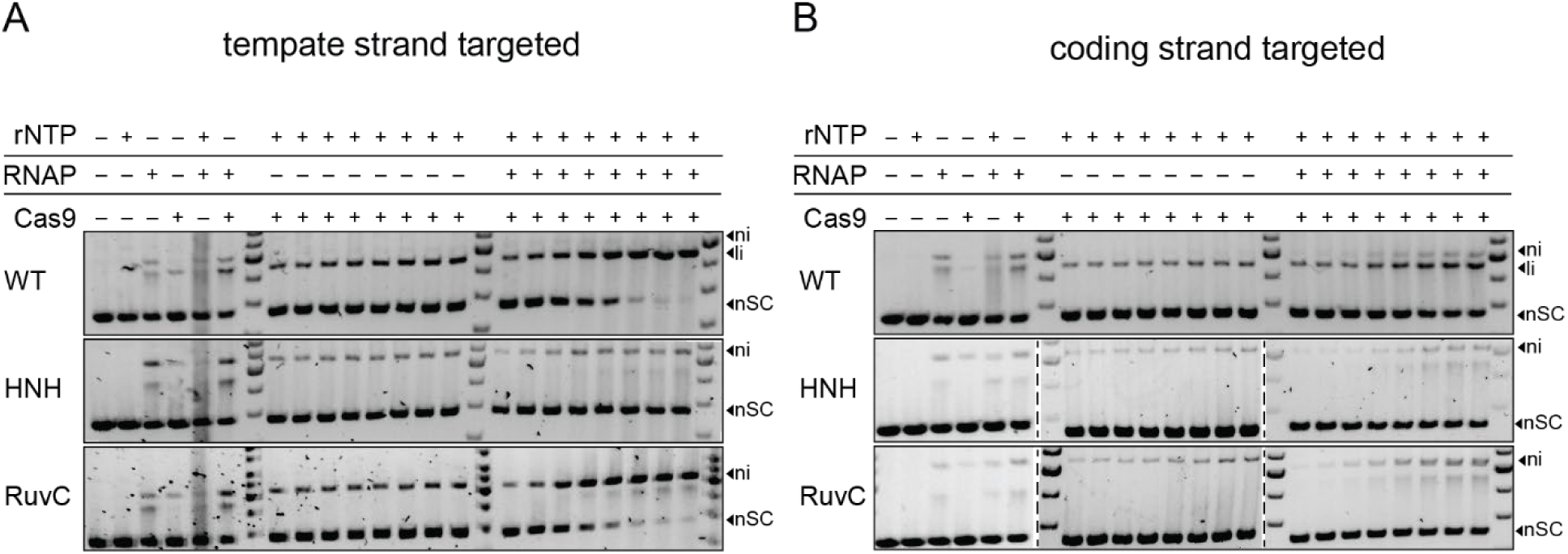
Gels showing Cas9 turnover by T7 RNA polymerase Cleavage assays were performed with an excess of plasmid DNA over Cas9. In the absence of RNAP, only a fraction of the DNA is cleaved due to the lack of Cas9 turnover. When RNAP is added in the presence of rNTPs, Cas9 can turnover under some conditions tested, resulting in full cleavage of the DNA at longer time points. A. The template strand of the target plasmid was targeted. B. The coding strand of the target plasmid was targeted.

## Supplementary Tables

List of λ genome targets

Primers for *cas9* mutagenesis and plasmids for Cas9 expression qPCR primers

Oligonucleotides for crRNA synthesis and targets for cleavage assays

